# Neuroblast migration along cellular substrates in the developing porcine brain

**DOI:** 10.1101/2021.08.19.456958

**Authors:** Demisha D.L. Porter, Sara N. Henry, Sadia Ahmed, Paul D. Morton

## Abstract

In the past decade it has become evident that neuroblasts continue to supply the human cortex with interneurons via unique migratory streams shortly following birth. Due to the size of the human brain, these newborn neurons must migrate long distances through complex cellular landscapes to reach their final locations. This process is poorly understood, largely due to technical difficulties in acquiring and studying neurotypical postmortem human samples along with diverging developmental features of well-studied mouse models. We reasoned that migratory streams of neuroblasts utilize cellular substrates, such as blood vessels, to guide their trek from the subventricular zone to distant cortical targets. Here we evaluate the association between young interneuronal migratory streams and their preferred cellular substrates in gyrencephalic piglets during the developmental equivalent of human birth, infancy, and toddlerhood.

## Introduction

The mammalian cortex varies in size and organization across species, but neurogenesis and neuronal migration has been evolutionarily preserved in both human and non-human primates; in particular, neocortical expansion has benefited from the continuous production of neurons throughout development into adulthood (Alvarez-Buylla and Garcia-Verdugo, 2002; Azevedo et al., 2009; Lois and Alvarez-Buylla, 1994; Luskin, 1993; Walton, 2012). Embryonically, the vast majority of neurons are generated in the germinal region lining the ventricle (the ventricular zone (VZ)), in which neural stem cells extend long processes to the surface wall of the cortex facilitating radial migration (Kriegstein and Noctor, 2004; Turrero Garcia and Harwell, 2017). During early postnatal periods of development a secondary proliferative area emerges, the subventricular zone (SVZ), housing the largest pool of neural stem/progenitor cells (NSPCs) further promoting cortical expansion (Sanai et al., 2005). These NSPCs give rise to immature neurons (neuroblasts) that migrate to their final destinations in various regions of the human brain, including the olfactory bulb (OB), the ventromedial prefrontal cortex (vPFC), and the anterior cingulate cortex (Aoyagi et al., 2018; Lazarini and Lledo, 2011; Lois et al., 1996; Paredes et al., 2016a; Sanai et al., 2011; Sawamoto et al., 2011). The journey from site of origin to final destination is largely influenced by the size and complexity of the brain, as neuroblasts must migrate significantly further through dense cellular networks in larger brains.

To supply the brain in such a manner requiring long migratory treks, it has been speculated that immature neurons utilize cellular substrates for guidance. Two main cellular substrates have been proposed to facilitate neuroblast migration, namely glial cells (gliophilic) and blood vessels (vasophilic). Within the postnatal and adult mammalian brain, elongated chains of migrating neuroblasts are tightly packed within glial tubes, encased by astrocytic processes, in the SVZ/RMS core (Gengatharan et al., 2016; Kornack and Rakic, 2001; Lois et al., 1996; Peretto et al., 2005). Impairments to the organization of glial tubes in postnatal and adult mice, were associated with chain migration defects to the OB, suggesting that neuroblast-astrocyte interactions are essential for this mode of transport (Anton et al., 2004; Belvindrah et al., 2007; Chazal et al., 2000; Kaneko et al., 2010).

The vascularization of the central nervous system is timed to coincide with the formation of the neuronal compartment during development (Paredes et al., 2018). Notably, blood vessels and their constituent endothelial cells support neuronal migration in several ways including (i) supplying oxygen and nutrients to meet the required metabolic demands of neuroblasts and (ii) providing physical scaffolds that secrete chemo-attractive/trophic factors, which ensures a specialized microenvironment to regulate neuroblasts migration, as well as maintenance (Fujioka et al., 2019; Karakatsani et al., 2019; Tata and Ruhrberg, 2018). Previous studies have demonstrated the migration of SVZ-derived neuroblasts along blood vessels tangentially and radially towards the cortex and OB giving rise to interneurons under normal physiological conditions and towards sites of injury to replace damaged neurons in cases of insults such as traumatic brain injury, stroke, and hypoxia-ischemia (Arvidsson et al., 2002; Bovetti et al., 2007; Christie and Turnley, 2012; Jinnou et al., 2018; Le Magueresse et al., 2012; Marin and Rubenstein, 2003; Metin et al., 2006; Parent et al., 2002; Snapyan et al., 2009).

Recent analyses of human postmortem tissue revealed unique migratory streams in neonates persisting through infancy: (i) medial migratory stream, and (ii) a collection of streams termed “Arc” supplying late-arriving interneurons to the frontal lobe, including the ventromedial prefrontal and cingulate cortices, respectively (Paredes et al., 2016a; Sanai et al., 2011). Yet the cellular processes enabling these unique migratory routes over vast distances remains elusive largely owing to difficulties in acquiring scarce neurotypical human postmortem tissue during these early epochs of life and technical difficulties limiting the number and diversity of experiments that can be performed; therefore, it is necessary to identify suitable animal model organisms as an alternative to address these questions.

Neonatal piglets represent a close bioequivalent to humans as they share similarities in gyrification, physiology, immunology, and anatomy. It was recently demonstrated that newborn piglet brains possess comparable migratory streams which provide the frontal lobe with newborn interneurons (Morton et al., 2017). As these cells are challenged with long migratory routes through a similar cytoarchitectural landscape to that of humans, we evaluated their potential embryonic origins, cortical destinations, and preferred cellular substrates from birth through toddlerhood in this report. We found that neuroblasts are closely associated with blood vessels and astrocytes during migration, as opposed to isolated chain-migration. Molecular profiling revealed that these migratory streams are largely comprised of immature neocortical interneurons derived from the MGE and CGE.

## Results

### Developmental timeline and study design

In recent years, larger mammalian species have emerged as ideal translational animal models for cell-dynamic studies of human brain development (Fernandez-Flores et al., 2018; La Rosa et al., 2018; Low et al., 2013; Parolisi et al., 2017). Piglets, unlike rodents, possess a highly evolved gyrencephalic neocortex (Fig. 1), large volume and similar distribution of white matter, and share key developmental trajectories similar to human postnatal brain development (Conrad et al., 2012; Costine et al., 2015). Recent reports showed that key neuronal niches lining the SVZ of the postnatal porcine brain resemble those reported in humans including cytoarchitectural lamination and robust migration of young interneurons following birth (Morton et al., 2017; Paredes et al., 2016a). In addition, numerous streams - seen as round clusters in orthogonal planes - of migrating neuroblasts were identified within the piglet SVZ oriented toward the cortex, as determined by expression of molecular markers DCX and PSA-NCAM (Morton et al., 2017). Thus, based on these characteristics we first evaluated the distribution of doublecortin (DCX^+^) neuroblast clusters in the SVZ at three stages of development: birth, infancy, and toddlerhood determined by longitudinal comparisons between pig and human brain growth (Fig. 1) (Conrad et al., 2012).

**Figure 1.**
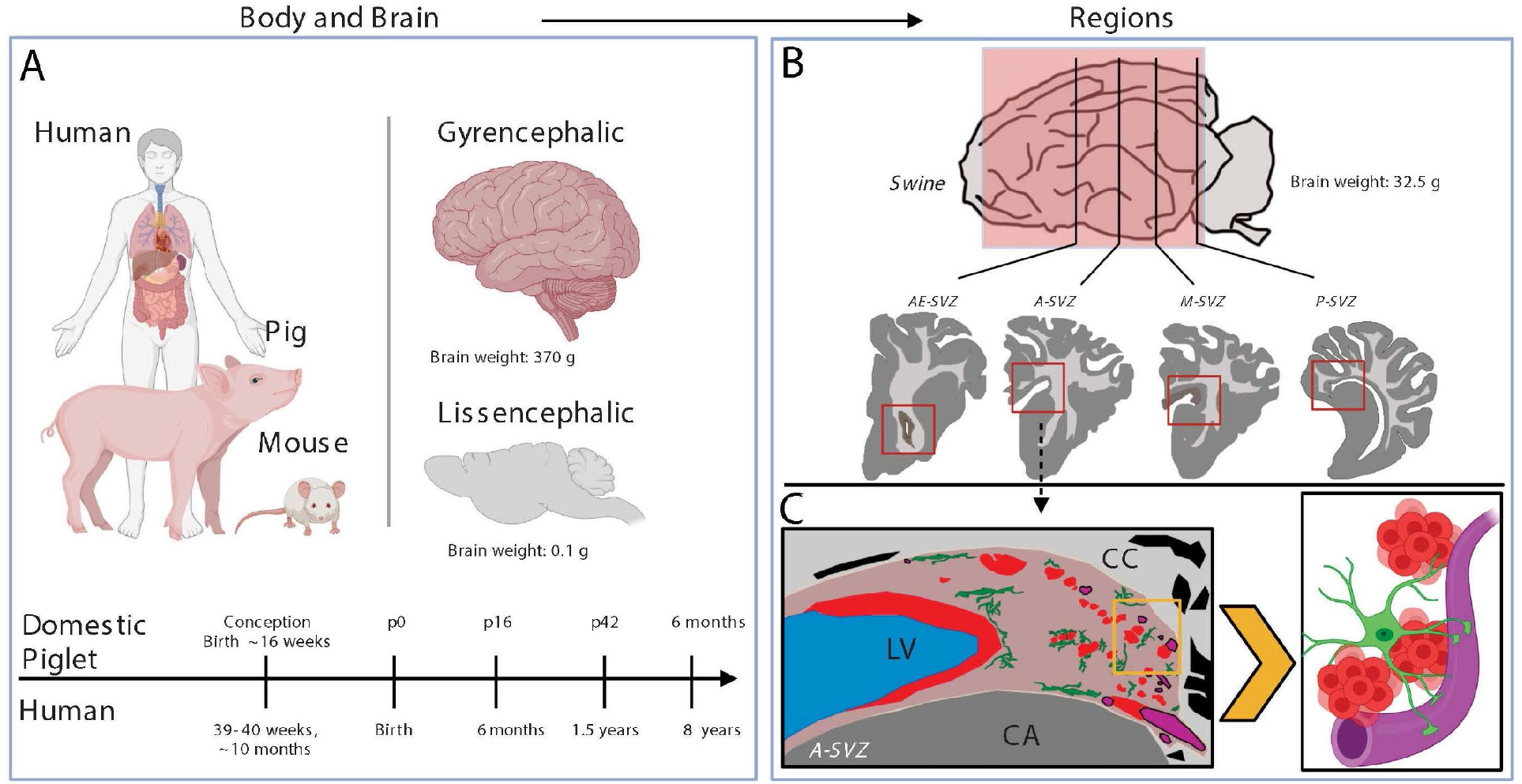
The domestic pig as a translational model for studying postnatal neurodevelopment. Comparison of human, pig, and mouse brain size, architecture, and developmental timeline (A). Brain weights are relative to a newborn period of life across species. (B) Illustration of the four rostro caudal brain regions (anterior end (AE-), anterior (A-), middle (M-), posterior (P-) subventricular zone (SVZ)) of interest in the whole porcine brain. (C) Cartoon illustration of the neuroanatomical and cellular composition of the A-SVZ. Neuroblasts (red); astrocytes (green); blood vessels (magenta). Abbreviations: CC corpus callosum; LV, lateral ventricle; CA, caudate nucleus. Illustrations presented in A and the right panel of C were generated using Biorender software.

### Spatiotemporal distribution and numbers of DCX^+^ neuroblast clusters in the postnatal porcine SVZ

To determine the spatiotemporal distribution of neuroblast clusters, the piglet SVZ was subdivided into four domains along the rostrocaudal axis to account for the regional differences in the human and porcine SVZ: the anterior end (AE)–, anterior (A)–, middle (M)–, and posterior (P)-SVZ (Fig. 2A-D)(Quinones-Hinojosa et al., 2007; Quinones-Hinojosa et al., 2006; Sanai et al., 2011). The boundary of the SVZ was easily recognizable based on previous reports, including distinct anatomical features: (i) opening of the lateral ventricle, (ii) lining of the corpus callosum dorsally, (iii) and head of the caudate nucleus inferiorly (Fig. 2G-R) (Costine et al., 2015; Morton et al., 2017). Histological screening revealed no significant differences in the total number of DCX^+^ neuroblast clusters between age groups throughout the entire SVZ (Fig. 2E); notably, DCX^+^ neuroblast clusters were in close proximity to IB4^+^ blood vessels in all regions and ages assessed (Fig. 2G-R). DCX^+^ neuroblast clusters were still detectable at lower levels at 6-months of age in the SVZ regions of the porcine brain (Fig. S1A-D); however, they appeared less compact and organized. These data suggest that young piglets generate abundant neuroblast clusters at birth which may persist into adulthood as described previously in some larger mammalian species (Parolisi et al., 2017; Piumatti et al., 2018).

**Figure 2.**
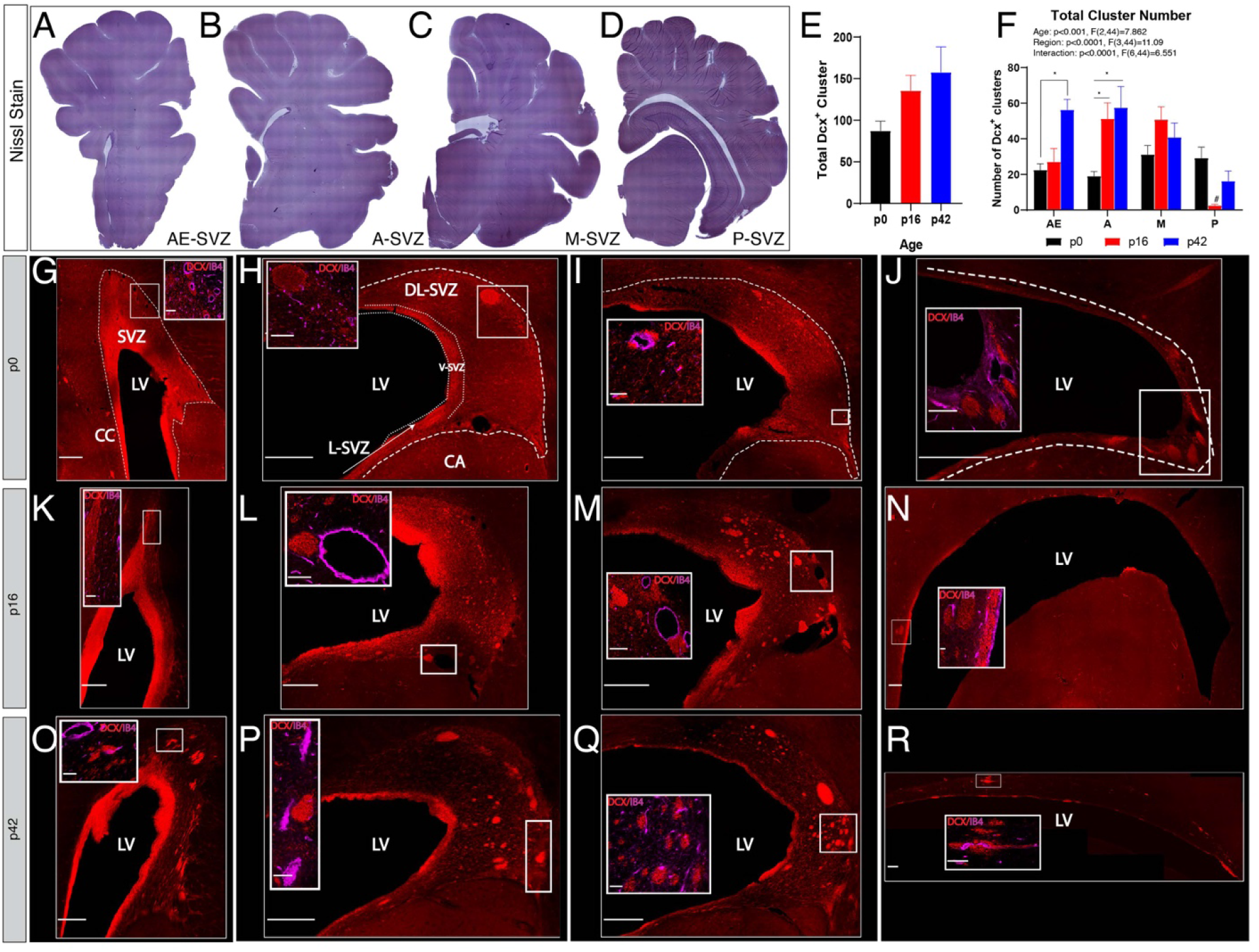
Development-related changes in the occurrence and regional distribution of neuroblast clusters throughout the piglet SVZ. (A-D) Cresyl violet staining on coronal sections of the SVZ at four anterior-posterior brain levels at 2 weeks of age. (E) Quantification of the number of DCX^+^ neuroblasts present in the SVZ (n=6 animals, p0; n=4 animals, p16; n=4 animals, p42). (F) Quantification of the number of DCX^+^ neuroblasts in the four SVZ subregions at p0, p16, and p42. (G-R) Spatiotemporal distribution of DCX^+^ neuroblasts in coronal sections; between birth and 42 days, DCX^+^ neuroblasts appear to have migrated from the posterior SVZ towards the rostral regions (A-SVZ) and in number with age. Higher magnification images of boxed areas show DCX^+^ neuroblasts close to IB4^+^ blood vessels in four SVZ rostrocaudal regions at p0, p16, and p42. Abbreviations: CC corpus callosum; LV, lateral ventricle; V-SVZ, ventricular-subventricular zone; SVZ, subventricular zone; DL-, dorsal-lateral; and L-, lateral. Data expressed as means ± SEM. One-way ANOVA with Bonferroni post hoc comparisons (E). *p<0.05, ^#^p<0.0001, analysis of variance (ANOVA) with Bonferroni post hoc test (F). Scale bar, 500 μm (insets, 100 μm).

Because the number of DCX^+^ neuroblast clusters appeared to increase with age, we next compared the occurrence and distribution of neuroblast clusters within the different SVZ subregions. Compared with newborns (p0), the number of DCX^+^ neuroblast clusters were significantly higher in animals at p42 in the AE-SVZ, and p16 and p42 in the A-SVZ (Fig. 2G). Two-way ANOVA analyses confirmed an interaction between age and subregion and DCX^+^ neuroblast clusters at p16 were significantly less abundant compared to the p0 and p42 in the P-SVZ (Fig. 2F). Collectively, these findings indicate that the piglet brain possesses a heterogeneous distribution of DCX^+^ neuroblast clusters in the SVZ, that are steadily maintained with age in a region-specific manner during postnatal porcine development. Because a collective majority of neuroblasts clusters were found within the A- and M-SVZ (Fig. 2F) and previous studies in piglets identified the A-SVZ as the most abundant source of newborn neurons capable of migrating to the frontal lobe at 2 weeks of age, we next restricted our analyses to the A-SVZ at p16.

### DCX^+^ neuroblast clusters express markers of migration in the porcine SVZ during postnatal development

To confirm that these clusters consist of migrating neuroblasts, we explored additional molecular markers. Polysialylated-neural cell adhesion molecule (PSA-NCAM) is expressed in migrating cells including neuroblasts within the SVZ (Bonfanti, 2006; Rutishauser, 2008). DCX^+^ neuroblast clusters co-expressed PSA-NCAM within the SVZ, which is indicative of migrating neuroblasts in the developing postnatal piglet brain (Fig. 3A-E). There were, however, some individual DCX^+^ cells within the neuroblast clusters that did not express PSA-NCAM. Therefore, we sought to determine if DCX^+^ PSA-NCAM^-^ neuroblasts clusters were undergoing proliferation or transitioning into mature neurons within the SVZ. We found that DCX^+^ neuroblast clusters rarely expressed the proliferative marker, Ki67 (Fig. 3F,G), or the mature neuronal marker, NeuN (Fig. 3H). Altogether, these data demonstrate that DCX^+^ neuroblast clusters in the postnatal SVZ are likely immature migrating neurons.

**Figure 3.**
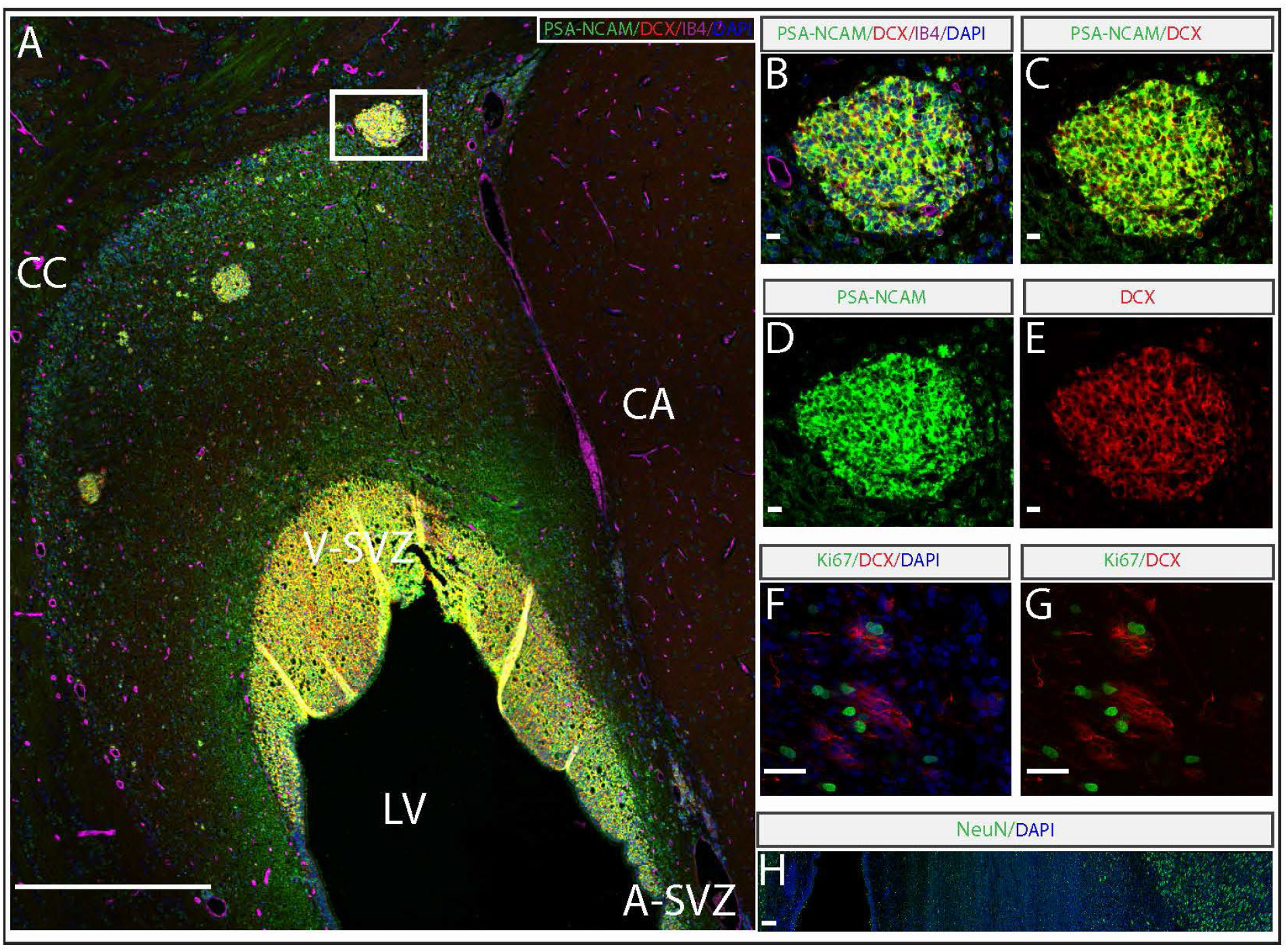
Postnatal neuroblast clusters express markers indicative of migration. Low magnification image (A) of a coronal section illustrating subpopulations of DCX^+^ neuroblasts in the A-SVZ illustrating co-expression with PSA-NCAM at 2 weeks of age; CC corpus callosum; CA, caudate nucleus; LV, lateral ventricle; V-SVZ, ventricular-subventricular zone; scale bar, 500 μm. (B-E) Higher magnification images of DCX^+^ PSA-NCAM^+^ neuroblasts, annotated by white box, DAPI, and IB4 immunostains; scale bars, 10 μm. (F-G) Piglet A-SVZ immunostained for cell proliferation marker, Ki67. Cell nuclei were stained with DAPI; scale bar, 100 μm. (H) Immunostain of DCX^+^ neuroblasts and the postmitotic neuronal marker NeuN; scale bar 100 μm.

### SVZ-derived migrating neuroblast clusters are largely comprised of young GABAergic interneurons in the postnatal piglet brain

Previous studies have shown that the neurogenic niches established during embryogenesis within the ventral subpallium, called the ganglionic eminences (GE), appear to be preserved until adulthood in the SVZ (Paredes et al., 2016a; Parnavelas et al., 2002; Rakic, 1988). Specifically, the spatial organization of progenitor cells within the GE, selectively express transcription factors that correspond to the production of distant neuronal subtypes. This led us to test whether the same applies to the DCX^+^ neuroblast clusters in the postnatal piglet SVZ. We found that DCX^+^ neuroblast clusters at p16 expressed the main inhibitory neurotransmitter *γ*-aminobutyric acid (GABA) an essential regulator of neocortical circuitry homeostatic balance (Fig. 4A-C). Since the GE can be further subdivided into the lateral ganglionic eminence (LGE), medial ganglionic eminence (MGE), and caudal ganglionic eminence (CGE), we focused our analyses on the cortical interneurons that primarily arise from the MGE and CGE(Ma et al., 2013; Nery et al., 2002). Within the A-SVZ at p16, neuroblast clusters expressed the calcium binding protein, secretagogin (SCGN)^+^ [associated with the CGE], which are selectively present in human cortical interneurons but not rodents(Raju et al., 2018). Additionally, individual cells within the clusters expressed markers of interneuron subclasses including parvalbumin and calretinin (Fig. 4D,E). Although not present within the neuroblast clusters, we identified a neuropeptide Y (NPY)-positive interneuron (Fig. 4F). To our surprise, we observed a small population of PV^+^CR^+^ cells within the clusters (Fig. 4G), an observation previously reported from other regions of large-brained gyrencephalic species, including humans (Hashemi et al., 2017; Leuba and Saini, 1997; Yan et al., 1995). In addition, all subtypes of interneurons assessed were clearly visible within the upper layers of the prefrontal cortex during an epoch of circuitry refinement (Fig. 4H,I).

**Figure 4.**
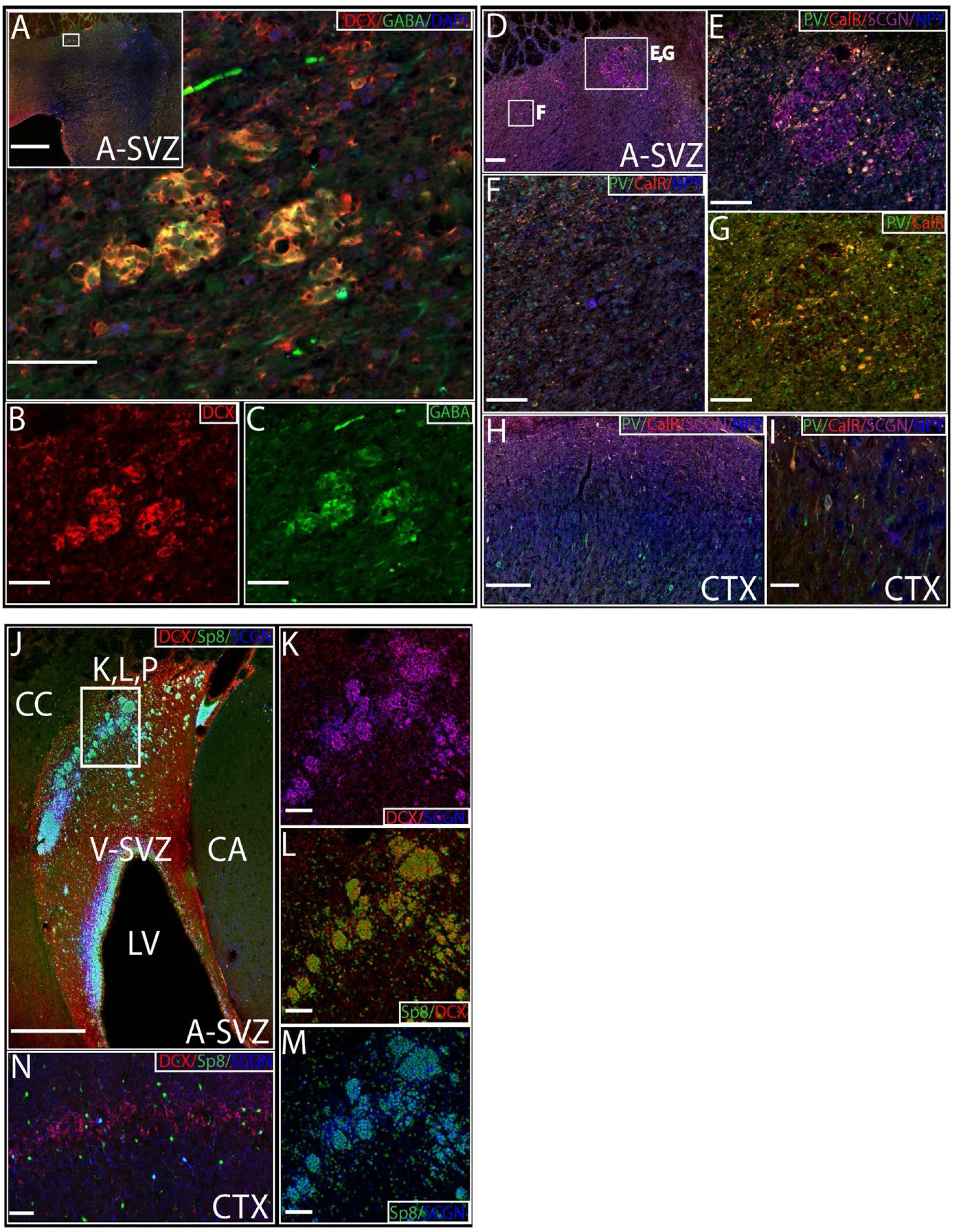
Detection of interneuron and subpallial marker expression in migrating neuroblasts within the SVZ of postnatal piglets. At 2-weeks of age, many of the migrating DCX^+^ neuroblasts in the A-SVZ express the inhibitory neurotransmitter GABA (A-C); scale bar 100 μm (inset, 500 μm). (D) Subpopulations of DCX^+^ neuroblasts/cells in the A-SVZ (regions noted in (E, F, G)) co-express interneuron subtype markers parvalbumin (PV), calretinin (CalR), or neuropeptide Y (NPY); scale bar 100 μm (D) 50 μm (E-G). PV^+^, CalR^+^, NPY^+^, and SCGN^+^ interneurons in the upper cortical layers I and II/III (H, I); scale bar 100 μm. (J) Individual DCX^+^ clusters express the regional transcription factor Sp8 specifying ventral telencephalic origin; scale bar, 500 μm. (K-M) DCX^+^ neuroblasts co-express SCGN and Sp8 in the A-SVZ; scale bar, 100 μm. Magnified images from the location marked by inset in (J). Individual cells express DCX^+^ SCGN^+^, and Sp8^+^ in the upper layers of the prefrontal cortex– (N). Abbreviations: CC corpus callosum; CA, caudate nucleus; LV, lateral ventricle; V-SVZ, ventricular-subventricular zone.

Reports in gyrencephalic species have shown co-expression of SCGN^+^ interneurons with Sp8, a marker associated with CGE/LGE origins (Ellis et al., 2019; Raju et al., 2018). Within the A-SVZ of p16 piglets there was robust co-expression of DCX^+^ neuroblast clusters with SCGN and Sp8 (Fig. 4J-M). DCX^+^Sp8^+^SCGN^+^ cells were also observed in the upper layers of the frontal cortex (Fig. 4N). Taken together, our results indicate that neuroblast clusters in the SVZ are young migrating GABAergic interneurons, likely derived from specific subregions of the ganglionic eminences that are destined to populate the postnatal neocortex.

### Migrating neuroblasts are closely associated with cellular substrates

In human infants, large numbers of neuroblasts have been documented in close association with astrocytes and blood vessels potentially serving as scaffolds to facilitate migration (Paredes et al., 2016a). Thus, we sought to determine whether SVZ-derived neuroblast clusters prefer the use of cellular substrates during normal development in the postnatal piglet. To visualize the association of cellular substrates near the surface of migrating neuroblast clusters, we generated a three-dimensional (3D) segmentation and rendering of immunolabeled clusters (Fig. 5A,B); axial image generated from 3D porcine atlas (Saikali et al., 2010). Within the dorsal-lateral SVZ, DCX^+^ neuroblast clusters were encased in GFAP^+^ astrocytic processes that appeared to be mostly aligned in the directionality (tangentially) of migration (Fig. 5C). In addition, neuroblasts clusters were observed directly contacting blood vessels in the A-SVZ, presumably utilizing them as a migratory substrate (Fig. 5D). Therefore, we quantified the distribution of DCX^+^ neuroblast clusters and measured the distance from blood vessels at all postnatal ages throughout the whole SVZ. Analysis of the average distance between clusters and vessels displayed a significant increase with age (Fig. 5E). We found that a majority of the DCX^+^ neuroblast clusters are within 0-20 µm distance of a blood vessel (average distance from the vessels for DCX^+^ neuroblast clusters: p0 = 5.13 µm, p16: = 12.80 µm, and p42 = 19.51 µm) (Fig. 5F). This suggests that as the brain develops, there is less association between streams of migratory neuroblasts and vascular substrates within the SVZ.

**Figure 5.**
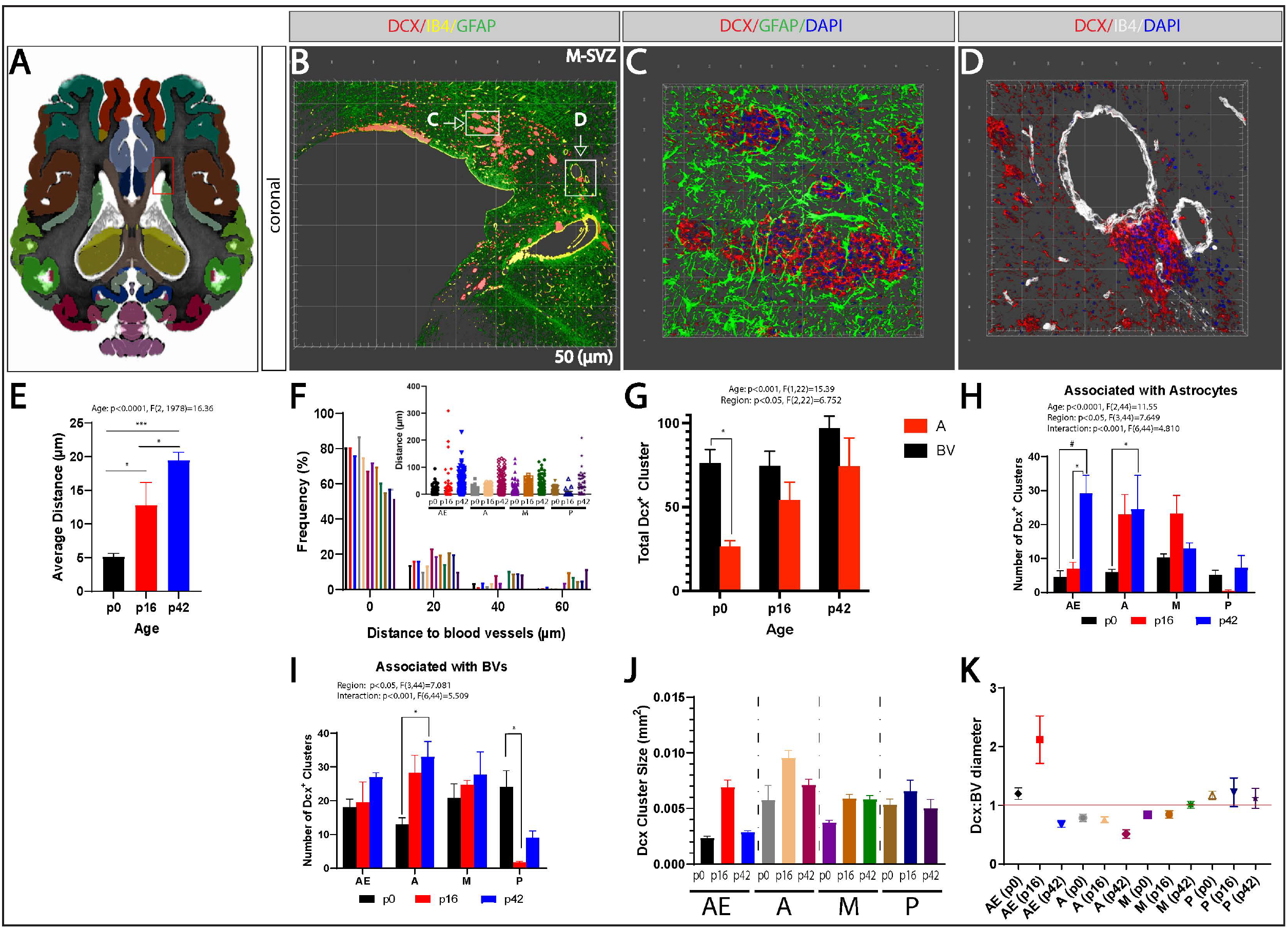
Migrating neuroblasts are commonly associated with cellular substrates in the SVZ of neonates. (A) Axial scan of pig brain from swine brain atlas; red box, location of A-SVZ. Cortices are color coded: red, dorsal prefrontal; dark green, primary somatosensory; brown, insular; light green, auditory; gray, middle temporal gyrus; burgundy, associative visual; forest green, parahippocampal; purple, cerebellum. (B) 3D-reconstructed image of the M-SVZ immunolabeled with DCX, IB4, and GFAP on the coronal plane. (C-D) 3D segmentation of DCX^+^ neuroblasts within the SVZ marked by insets in (B), illustrating DCX^+^ neuroblasts encased in GFAP^+^ astrocytic processes (C), and intimately associated with IB4^+^ blood vessels (D). (E) Quantification of the average distance of between DCX^+^ clusters and blood vessels. (F) Frequency distribution of DCX^+^ neuroblast cluster-to-vessel distance measured from the edge of each cluster to its nearest neighboring the entire SVZ. Inset shows the quantitated average of cluster to vessel distance within SVZ subregions. (G) Quantification of the number of DCX^+^ clusters in association with GFAP^+^ astrocytic processes or IB4^+^ blood vessels between birth and 42 days of age. (H-I) Quantification of the number of clusters associated with astrocytes (H) or blood vessels (I). (J) Size of DCX^+^ neuroblasts measured throughout the entire SVZ at different ages. (K) Size correlation between blood vessel and DCX^+^ neuroblast cluster diameter. Data is expressed as the mean ±SEM. n=4-6 animals/group. One-way ANOVA with Bonferroni post-hoc comparisons (E). *p<0.05, #p<0.001, ***p<0.0001, two-way ANOVA with Bonferroni post-hoc comparisons (G-J).

Next, we quantified the association of DCX^+^ neuroblast clusters with cellular substrates across age and SVZ subregions. Quantification of the number of neuroblast clusters associated with astrocytes and blood vessels at all ages, revealed that their number and association with astrocytes was significantly higher at birth compared to blood vessels at all ages, p0-p42 (Fig. 5G). However, there were no significant differences in their association with blood vessels across ages. To further investigate the occurrence and distribution of neuroblast clusters with cellular substrates, we analyzed the different SVZ subregions during postnatal development. Here, we found a dramatic and significant increase in the number of neuroblasts associated with astrocytes across development in the AE-SVZ. In addition, this was observed within the A-SVZ region at p0 compared to p42 (Fig. 5H), suggesting that these chains of neuroblast clusters seem to be directed towards the olfactory bulb through the glial tubes as this sensory modality develops.

Similarly, we found that the DCX^+^ neuroblast clusters showed a significant increase in association with blood vessels within the A-SVZ from p0 to p42. However, in the P-SVZ, at p0 there was a significant reduction in the amount of DCX^+^ neuroblast clusters associated with blood vessels at p16 (Fig. 5I). DCX^+^ neuroblast clusters appeared heterogeneous in size (ranging from .0024 to .0096 mm^2^, in cross-sections) (Fig. 5J). Moreover, when we assessed the relationship of DCX^+^ neuroblast cluster diameter expressed as a ratio to their neighboring blood vessel diameter, the majority of the clusters in the A- and M-SVZ across ages appeared to be smaller than the blood vessels (Fig. 5K). From this data, we concluded that the extent of neuroblast association with glial and blood vessel substrates corresponded to the postnatal changes in distribution of neuroblast clusters within the different SVZ subregions (see Fig. 2).

### Postnatal expression of SCGN^+^Sp8^+^DCX^+^ neuroblast clusters associated with blood vessels in the porcine SVZ

Due to the wide expression of SCGN and Sp8 throughout the porcine SVZ in DCX^+^ neuroblast clusters, we investigated whether SCGN^+^Sp8^+^ clusters associated with blood vessels. In the A-SVZ of p0 and p16 piglets, each DCX^+^ cluster co-expressed SCGN^+^Sp8^+^ (Fig. 6A-E). At birth, the percentage of co-localization between DCX and SCGN varied to that seen in DCX^+^Sp8^+^ clusters. The association of DCX^+^SCGN^+^ clusters with blood vessels appeared to be similar throughout development, compared to DCX^+^Sp8^+^ cluster. However, the percentage of co-localized clusters increased with age in both groups. Moreover, the percentage of migrating DCX^+^SCGN^+^ clusters that were not in contact with blood vessels was almost identical with DCX^+^Sp8^+^ clusters during the early postnatal period (Fig. 6F-G). These findings suggest that during critical periods of postnatal development, cortical GABAergic interneurons preferably associate with blood vessels for neuronal migration to distant locations within the brain.

**Figure 6.**
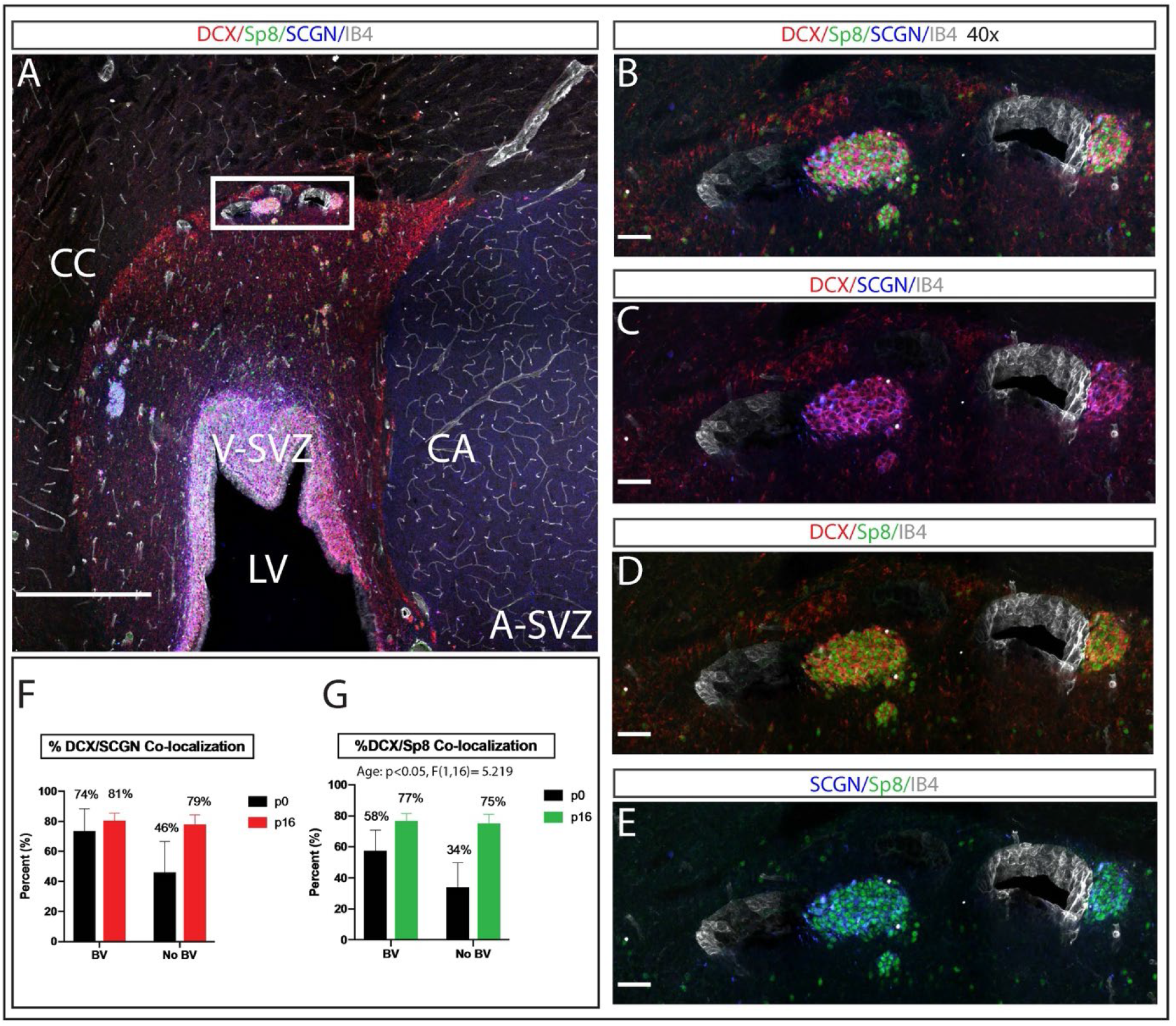
A majority of DCX^+^ neuroblast clusters co-localize with SCGN and Sp8 in the piglet SVZ. (A) DCX, Sp8, SCGN, IB4 immunolabeled coronal section at 2-weeks of age, illustrating the contact between neuroblast clusters and blood vessels in the A-SVZ; scale bar, 500 μm. Higher magnification images of DCX^+^Sp8^+^SCGN^+^ clusters (B-E); scale bar, 100 μm. Quantification of co-localization with SCGN (F) or Sp8 (G) in the presence and absence of vessel association. Data is expressed as the mean ±SEM. n=4-6 animals/group. Two-way ANOVA with Bonferroni post-hoc test (G). Abbreviations: CC corpus callosum; CA, caudate nucleus; LV, lateral ventricle; V-SVZ, ventricular-subventricular zone.

## Discussion

In the present study, we profiled streams of neuroblasts spatially and temporally throughout the porcine SVZ during early brain development. We revealed that DCX^+^ neuronal clusters are present primarily in the anatomical equivalent of tier III of the human SVZ and are maintained throughout early postnatal development. In contrast to studies in ferrets, monkeys, and humans that reported a distinct decline of DCX expression, we found that the number of DCX^+^ clusters remained constant from birth through toddlerhood in the piglet SVZ (Fig. 2) (Akter et al., 2020; Ellis et al., 2019; Paredes et al., 2016a; Sanai et al., 2011). These observations are in sharp contrast to previous findings in the piglet where DCX^+^ cells decline with age (birth to 15 wks) and human studies were migrating immature neurons (DCX^+^ cells) decline by 6 to 18 months of age (Morton et al., 2017; Paredes et al., 2016a; Sanai et al., 2011). However, previous studies evaluated individual cells, whereas these findings are focused exclusively on clusters of neuroblasts purported to be members of organized migratory streams.

The maintenance of DCX^+^ clusters could represent the retention of juvenile “non-newly generated cells” that are born prenatally and continuously express molecular markers of immature cells well into postnatal development as has been observed in layer 2 of the cerebral cortex of the sheep, cat, guinea pig, and reptiles (Cai et al., 2009; Luzzati et al., 2009; Piumatti et al., 2018; Xiong et al., 2008). In agreeance with the findings of developmental markers for structural plasticity and neuronal migration that transcend typical age-related expression patterns, immunohistochemical analysis revealed that the vast majority of DCX^+^ cells in the neuroblast clusters of the postnatal porcine SVZ expressed the migratory marker PSA-NCAM (Fig. 3, A-E), but did not express Ki67(Fig. 3, CC, D or NeuN (Fig. 3, E). Considering that Ki67 and NeuN are necessary for cell proliferation and neuronal maturation, respectively, it appears that porcine SVZ-derived neuroblasts are young migratory neurons. Our observations in conjunction with evidence from preserved subpopulations of immature neurons throughout various cortical regions, white matter, and SVZ-lr of cetaceans and mammals, implies that in the porcine SVZ, late-waves of young migratory neurons may be the source for new neurons that integrate into the existing cortical circuitry during postnatal life (La Rosa et al., 2018; Parolisi et al., 2017; Piumatti et al., 2018). Indeed, previous findings from focally labeled cells within the porcine A-SVZ demonstrated interneurons within the upper layers of the frontal lobe (Morton et al., 2017).

Within the postnatal and adult mammalian brain, neuroblasts in the SVZ have been reported to migrate to the frontal cortices, where their postmitotic features are associated with the expression of transcription factors derived from the embryonic ganglionic eminence (Cai et al., 2009; Ellis et al., 2019; Paredes et al., 2016a; Xiong et al., 2008). Using birth dating markers to determine the embryonic origin and possible fate of these DCX^+^ clusters, we discovered that neuroblasts within the SVZ are a heterogeneous population of GABAergic interneurons. Interestingly, we also found cells within the clusters, co-expressed the marker associated with the MGE, parvalbumin (PV^+^), and calretinin (CR^+^) [possibly associated with LGE] (Fig. 4 D-I), a phenomenon previously reported in the developing human cortex and the *Macaca* primary visual cortex (Hashemi et al., 2017; Leuba and Saini, 1997; Yan et al., 1995). This observation raises the possibility that within the developing gyrencephalic brain, the transient co-localization of PV^+^ and CR^+^ calcium-binding proteins is related to the structural/functional alterations associated with cortical plasticity leading into later stages of development. Furthermore, subsequent analyses revealed that the SVZ-derived DCX^+^ clusters were also Sp8^+/^SCGN^+^, two markers of interneurons originating from the CGE/LGE (Fig. 4 J-N). It has recently been shown that SCGN^+^ interneurons populate the upper and lower cortical layers of human and ferret brain tissue but are absent from the mouse cortex (Ellis et al., 2019; Raju et al., 2018). Hence, our study shows that the vast majority of the SVZ-derived DCX^+^ clusters originate in the subpallial region of the telencephalon and migrate to the neocortex, which is likely to contribute to cortical plasticity in the postnatal piglet. Further experimental analysis is needed to determine the fate of postnatal interneurons in the porcine model.

On account of the variability in brain size and cortical folding, the orders of magnitude for migration distances can vary between species depending on neurogenic origin and their final cortical destinations from infancy to adulthood (see review (Paredes et al., 2016b)). In this context, most SVZ neuroblasts must transverse through a densely compacted extracellular environment to reach their cortical targets. Here, we have shown that neuroblasts in the SVZ are encased by astrocytic processes and preferentially associated with blood vessels (Fig. 5 A-D). On the other hand, by p42, the number of neuroblasts encased by astrocytic processes was drastically increased in the rostral regions of the SVZ (Fig. 5 E-I). The apparent increase in neuroblast-astrocyte interaction is similar to studies in postnatal mice and rats, in which the assembly of glial tubes and chain migration from the SVZ to the OB was detected by p21-25 (Bozoyan et al., 2012; Peretto et al., 2005). Nevertheless, it’s become more and more evident that SVZ-derived neuroblasts utilize vascular scaffolds not only for migration within the RMS-OB but also through the parenchyma, as seen in postnatal rabbits; cerebral cortex as seen in mice; the SVZ arc as seen in human infants (Bovetti et al., 2007; Inta et al., 2008; Le Magueresse et al., 2012; Paredes et al., 2016a; Ponti et al., 2006; Snapyan et al., 2009). Moreover, our results showed that not all clusters’ sizes were equivalent to the same blood vessels, suggesting that cluster-blood vessel interactions are heterogeneous in the developing neocortex and vessels provide specialized microenvironments for these clusters via the delivery of nutrients and oxygen (Fig. 5 J, K); however, it is unclear whether the heterogeneities in size are stochastic. Taking into account the close relationship between DCX^+^ clusters and blood vessels in the developing human fetal brain, proximity to blood vessels may exacerbate vulnerability to pathological changes in blood flow, hypoxia, or hypoxia-ischemia, which would likely impact excitatory/inhibitory balance during the later stages of cortical growth.

The dynamic relationship between extended postnatal development and neocortical expansion remains to be understood (Stepien et al., 2021). Nevertheless, the overall distribution of postnatal DCX^+^ neuroblasts might represent a large, heterogeneous population of undifferentiated neurons that serves as a parallel form of plasticity in large-brained mammals. This hypothesis is supported by the persistent occurrence of undifferentiated cells, postnatally, with morphology and cell marker expression patterns reminiscent of immaturity in larger-expanded neocortices (Parolisi et al., 2017; Piumatti et al., 2018). Moreover, the migration modalities revealed here represent anatomical substrates which we infer SVZ-derived neuroblast clusters utilize for migration to the cortex in a substrate-dependent manner. Preferential association with the vascular system might also be important in terms of cortical maturation as the vascular niche is a critical regulator of cellular function. Defects to neuron migration along blood vessels could result in immature cortical development, as implied in a swine model of congenital heart disease (Morton et al., 2017). Future studies focused on substrate migration from the postnatal SVZ in large-animals, may yield novel complex migratory streams that are yet to be discovered in the developing human brain.

### Experimental procedures

#### Animals

Domestic, White (Yorkshire/Landrace) piglets were obtained from the Swine Agricultural Research and Extension Center at Virginia Tech. All experiments were conducted in accordance with the NIH Guide for the Care and Use of Laboratory Animals and performed under the approval of the Virginia Tech Institutional Animal Care and Use Committee.

#### Tissue collection and processing

Quickly following chemical euthanasia, brains were removed from the skull, cut at 5mm intervals, and fixed at 4ºC in 4% paraformaldehyde (0.1 M PBS) for 72 h. After fixation, tissue slabs were cryoprotected at 4ºC in a sucrose gradient of 20% and 30% (0.1M PBS) until sunken. Tissue was embedded in OCT compound (Tissue-Plus #4585) and stored at −80°C for serial sectioning on a Thermo Scientific CryoStar NX50 cryostat. Coronal, sagittal, and horizontal tissue sections (50 μm) were collected for immunohistochemical procedures. For Cresyl Violet staining, 5mm tissue blocks were incubated in 10% formalin for 72 h, embedded in paraffin, and cut on a microtome at 5 μm thickness.

### Histology

#### Cresyl Violet staining

Paraffin-embedded sections (5 µm thick) were treated with xylene for 10 min and rehydrated in a descending series of ethanol steps followed by PBS: 100% 3×3 min, 95% 2×3 min, 70% 2×3 min, and PBS 2×10 min. Sections then underwent antigen retrieval at 95°C in 10 mM Na Citrate buffer, pH=6.0. Following antigen retrieval, slides were treated with 10% formalin for 10 min and 2% cresyl violet (Electron Microscopy Science, Hatfield, PA, USA) solution for 2hrs. Sections were rinsed in PBS for 5 min and coverslipped using SouthernBiotech mounting medium.

#### Free-floating immunohistochemistry

Primary antibodies that did not require tyramide signal amplification were stained in the following manner. Free-floating sections (50 μm) were incubated in blocking solution (20% normal goat or donkey serum in 0.1 M PBS) for 1.5 h at room temperature. Sections were then incubated in primary antibodies, diluted in carrier solution (1% bovine serum albumin, 1% normal goat or donkey serum, and 0.3% Triton X in 0.1 M PBS), overnight on an orbital shaker at 4ºC. Sections were washed 3 × 30 min each in PBS-T (0.2% Triton X-100 in PBS) on a rocker at room temperature. Sections were then incubated for 1.5 hr with species-specific, secondary fluorescent antibodies diluted in carrier solution (1:500). Sections were washed 3 × 30 mins each in PBS-T at room temperature. Sections were covered with coverslips using DAPI Fluoromount-G.

#### Tyramide signal amplification (TSA)

Free-floating sections (50 μm) were first mounted on glass slides and allowed to dry for 1 hr. Some antigens required antigen retrieval (see Table S1), which was conducted at 95°C in 10 mM Na Citrate buffer, pH=6.0. Following antigen retrieval, slides were washed with PBS-T for 10 min, placed in 1% H_2_O_2_ in PBS for 45 minutes, and then blocked with TNB solution (0.1 M Tris-HCl, pH 7.5, 0.15 M NaCl, 0.5% BSA) for 1 hour. Slides were incubated in primary antibodies overnight at 4°C (see Table S1) followed by 3 × 30 min rinses in PBS-T. Sections were then incubated in species-specific Biotin-SP-conjugated secondary (1:200; Jackson Immunoresearch Laboratories) for 2.5 hours at room temperature. All antibodies were diluted in TNB solution. Following 3 × 30 min rinses in PBS-T, sections were incubated in streptavidin-horseradish peroxidase for 30 min at room temperature. Sections were rinsed and incubated in a tyramide working solution (Invitrogen™) for 5 min at room temperature using the following dilutions: CF405S 1:200; Fluorescein, 1:100; Cy3, 1:200; Cy5, 1:200. Sections were then rinsed in PBS-T (1×30 min) and mounted with SouthernBiotech mounting medium.

### Imaging

A Zeiss LSM 880 (Carl Zeiss Microscopy) or Nikon C2 confocal laser scanning microscopic system (Nikon Instrument Inc., NY) was used in collecting tiled and high-resolution z-stacks for analyses following fluorescent immunohistochemical staining. Images were acquired with a 4x,10x, 20x, or 40x water objective. All images were acquired as 50μm thick sections. .nd2 and .czi files were covered to .sis files using Vision 4D software for further analyses. White light images were acquired at 4x on an Olympus BX51TRF microscope with MicroBrightField Stereoinvestigator software on 5 µm thick sections.

### Cell counting and quantification

#### Neuroblast cluster measurements

Exhaustive quantification of the number of DCX^+^ clusters was analyzed on tiled z-stack projections spanning two levels of the SVZ (dorsolateral (DL-) and lateral (L-) SVZ) using Vision 4D software. The size of each cluster was determined by manually drawing polygonal contours and calculating the area (µm); approximations of cluster diameter were further calculated based on the area of a circle.

#### Neuroblast cluster analyses relative to the cellular substrate

Quantification of DCX^+^ clusters near blood vessels and within astrocytic glial tubes, was performed using Vision 4D software. In each SVZ area, the total number of DCX^+^ clusters nearest to the blood vessels along with those surrounded by astrocytic processes were counted using Vision 4D software. The distance from each DCX^+^ cluster to the nearest blood vessel was manually calculated by an experienced researcher using the ruler annotation function of Vision 4D software. Distances in micrometers were recorded, and data are presented as a histogram of the frequency of clusters relative to the distance to the substrate.

#### Neuroblast cluster co-localizations

Confocal images were taken from coronal slices (50 μm thickness) within the anterior (A-) SVZ at 0, 16, and 42 days of age. Percent co-localizations were calculated by counting the number of co-localized cells and dividing by the total number of DCX^+^ cell bodies per cluster using ImageJ software.

### Statistical analyses

A two-tailed, unpaired Student’s t-test was performed for single comparisons; data were considered significantly different if p < 0.05. When two or more samples were being compared, statistical significance between groups was determined by two-way analysis of variance (ANOVA) with Bonferroni’s post hoc multiple comparison test; p < 0.05 determined significance. Data are expressed as mean ± SEM (standard error of mean). GraphPad Prism software was used to run Student’s t-test and ANOVAs. Error bars represent SEM, ^*^p ≤ 0.05, ^***^p ≤ 0.001, ^#^p ≤ 0.0001.

## Acknowledgments

We gratefully thank Amy Rizzo for her assistance with veterinary services. We would like to acknowledge Karen Hall and Rachel McNeil for animal care. This work was supported by the US National Institute of Health (R15NS108183 to PDM).

## Author contributions

PDM designed the experiments. DDLP carried out experiments with assistance from SNH and SA. DDLP analyzed data. DDLP prepared the figures and wrote the report with P.D.M.

## Data availability

All data are included in the manuscript and supplementary materials.

