## Supplementary Data for "Neuroblast migration along cellular substrates in the developing porcine brain"

**Supplemental information**

**Figure S1**

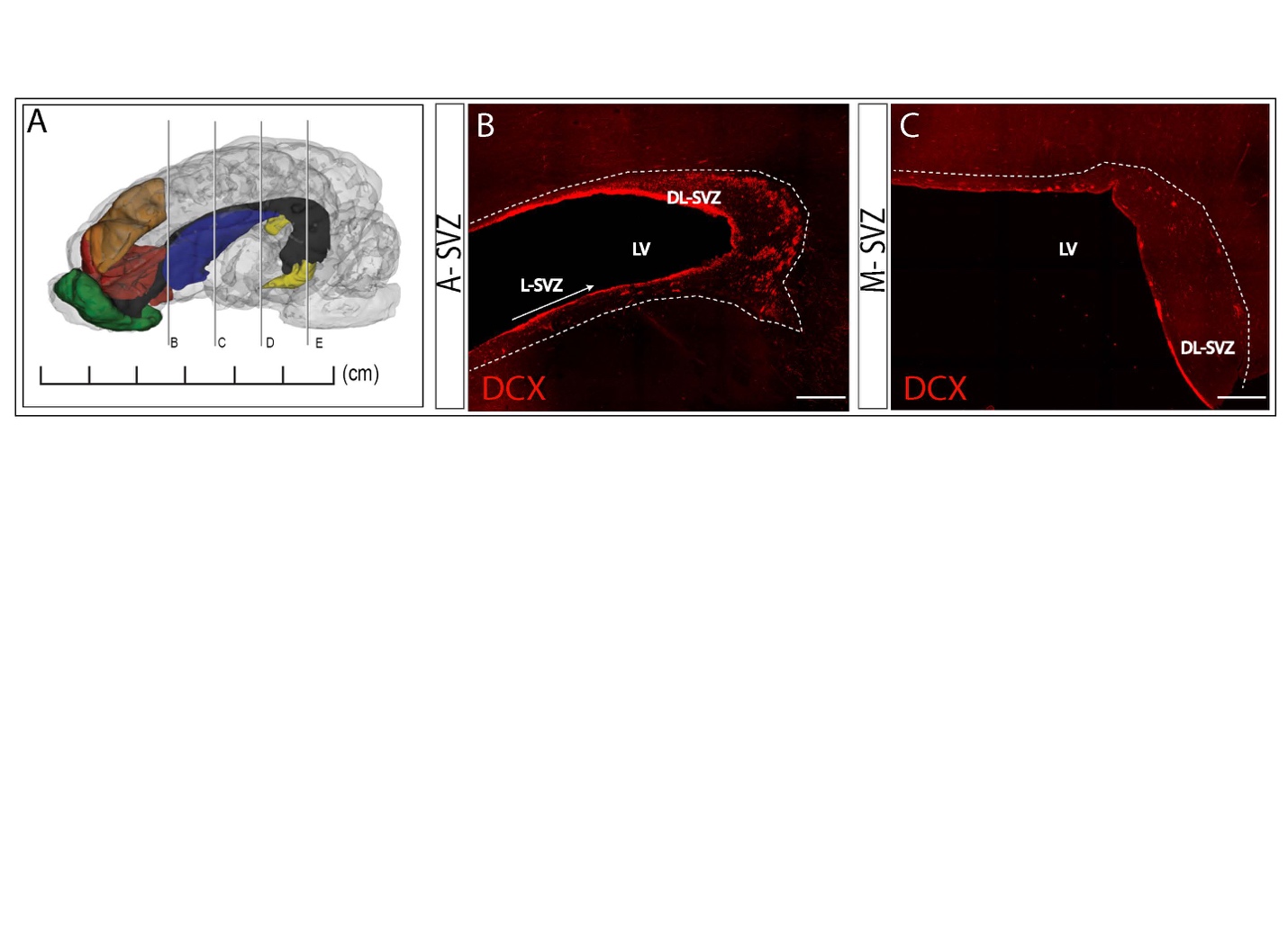

**Figure S1. Spatiotemporal distribution of DCX^+^ clusters in the 6-month piglet brain.** (A) 3D representation of the pig brain. Green, olfactory bulb; red, ventral prefrontal cortex; orange, dorsal prefrontal cortex; blue, caudate nucleus; black, lateral ventricle; yellow, hippocampus. (B,C) Dense collection of DCX^+^ clusters observed in coronal sections of the anterior (B) and middle (C) subventricular zone; scale bar 500 μm. DCX, doublecortin; DL, dorsolateral; L, lateral; LV, lateral ventricle; SVZ, subventricular zone.

**Table S1**

| Antibody | Catalog # | Vendor | Polyclonal/  Monoclonal | Host Species | Isotype | IHC Working Dilution (Pig) |
| --- | --- | --- | --- | --- | --- | --- |
| ^#^Calretinin | ABN2191 | EMD Millipore | Polyclonal | Rabbit | IgG | 1:200, AR 10min |
| *Doublecortin  (DCX) | ab18723 | Abcam | Polyclonal | Rabbit | IgG | 1:500, AR 5min |
| ^#^Doublecortin  (DCX) | ab2253 | EMD Millipore | Polyclonal | Guinea Pig | IgG | 1:500, AR 5min |
| ^#^Doublecortin  (DCX) | sc-271390 | Santa Cruz | Monoclonal | Mouse | IgG1 | 1:500, AR 10min |
| ^#^ gamma-  Aminobutyric acid (GABA) | A0310 | Sigma | Monoclonal | Mouse | IgG1 | 1:500, AR 5min |
| *Glial Fibrillary acidic protien  (GFAP) | ab4674 | Abcam | Polyclonal | Chicken | IgY | 1:500 |
| *Ki67 | 550609 | BD Pharmingen | Monoclonal | Mouse | IgG1,K | 1:50 |
| *NeuN | ABN91 | EMD Millipore | Polyclonal | Chicken | IgY | 1:500 |
| ^#^Neuropeptide Y | 22940 | ImmunStar | Polyclonal | Rabbit | IgG | 1:200, AR 10min |
| ^#^Parvalbumin | MAB1572 | EMD Millipore | Monoclonal | Mouse | IgG1 | 1:500, AR 8min |
| ^#^PSA-NCAM | MAB5324 | EMD Millipore | Monoclonal | Mouse | IgM | 1:500, AR 11min |
| ^#^SCGN | HPA006641 | Sigma | Polyclonal | Rabbit | n/a | 1:1,000, None |
| ^#^SP8 | HPA054006 | Sigma | Polyclonal | Rabbit | n/a | 1:200, AR 10min |
| *Isolectin GS-IB4 | I32450 | Thermofisher Scientific | n/a | n/a | n/a | 1:500 |

**Table S1. Antibodies used.** AR, antigen retrieval. *Free floating immunohistochemistry. ^#^Tyramide signal amplification.
